# Structure of the ISW1a complex bound to the dinucleosome

**DOI:** 10.1101/2023.01.02.522444

**Authors:** Lifei Li, Kangjing Chen, Youyang Sia, Pengjing Hu, Youpi Ye, Zhucheng Chen

**Author notes:** Correspondence to Zhucheng Chen.

## Abstract

Nucleosomes are basic repeating units of chromatin, and form regularly spaced arrays in cells. Chromatin remodelers alter the positions of nucleosomes, and are vital in regulating chromatin organization and gene expression. Here we report the cryoEM structure of chromatin remodeler ISW1a complex bound to the dinucleosome. Each subunit of the complex recognizes a different nucleosome. The motor subunit binds to the mobile nucleosome and recognizes the acidic patch through two arginine residues, and the DNA-binding module interacts with the entry DNA at the nucleosome edge. This nucleosome-binding mode provides the structural basis for linker DNA sensing of the motor. Notably, the Ioc3 subunit recognizes the disk face of the adjacent nucleosome through the H4 tail, the acidic patch and the nucleosomal DNA, which is important for the spacing activity in vitro, and for nucleosome organization and cell fitness in vivo. Together, these findings support the nucleosome spacing activity of ISW1a, and add a new mode of nucleosome remodeling in the context of a chromatin environment.

---

DNA wraps around the histone octamers by ~147 bp to form nucleosomes. The deposition of nucleosomes along the genomic DNA shows prominent features^1^.The promoter regions around the transcription start sites of active genes form nucleosome-free regions (NFRs), which are followed by well-positioned nucleosome arrays downstream ^2^. In contrast, in the repressed chromatin region, nucleosomes form evenly spaced arrays^3^, which largely occlude the access to the genetic materials. These features of chromatin organization are conserved from yeast to human cells, and important for regulation of gene transcription^4,5^.

Chromatin remodelers are ATP-dependent motor proteins that play major roles in nucleosome organization inside the cell^6,7^. Although the fundamental mechanism of chromatin remodelers acting on an individual nucleosome is extensively studied^8,9^, how they work in the context of nucleosome arrays is largely unexplored.

The ISWI family chromatin remodelers are responsible for the assembly of the nucleosome arrays inside the cell, and generally function in promoting repressed chromatin and gene silencing^10–12^. Isw1 is a key chromatin remodeler that controls nucleosome organization in yeast^13–16^. Isw1 interacts with Ioc3 to form the ISW1a complex^17^, which is particularly enriched at the +1 nucleosome, slides the nucleosome toward the upstream of the NFR, and represses the expression of many genes in yeast^13,15,18–20^. Isw1 also forms a ISW1b complex, which shifts the nucleosomes to the opposite direction inside the cells^20,21^. ISW1a shows a potent nucleosome spacing activity, setting up an inter-nucleosome linker DNA length of ~30 bp in vitro^16,18,22–24^. ISW1a shows a preference for the dinucleosome over mononucleosome in chromatin remodeling^25,26^.

Isw1 contains multiple functional domains ^27–29^, including the N-terminal AutoN domain, the central catalytic motor domain, the C-terminal inhibitory NegC domain, the polybasic arginine anchor (RA) motif, and the DNA-binding HAND-SANT-SLIDE (HSS) domain (Fig. 1a). Previous studies based on the structures of the DNA-binding module of ISW1a lacking the ATPase motor domain (HSS-Ioc3) suggested a protein ruler model in control of nucleosome spacing^17^. Recent studies revealed the interaction between the Isw1 ATPase motor and the mononucleosome^30–32^. It remains unknown how the ISW1a complex functions as a unit to control the promoter nucleosome organization.

**Figure 1.**
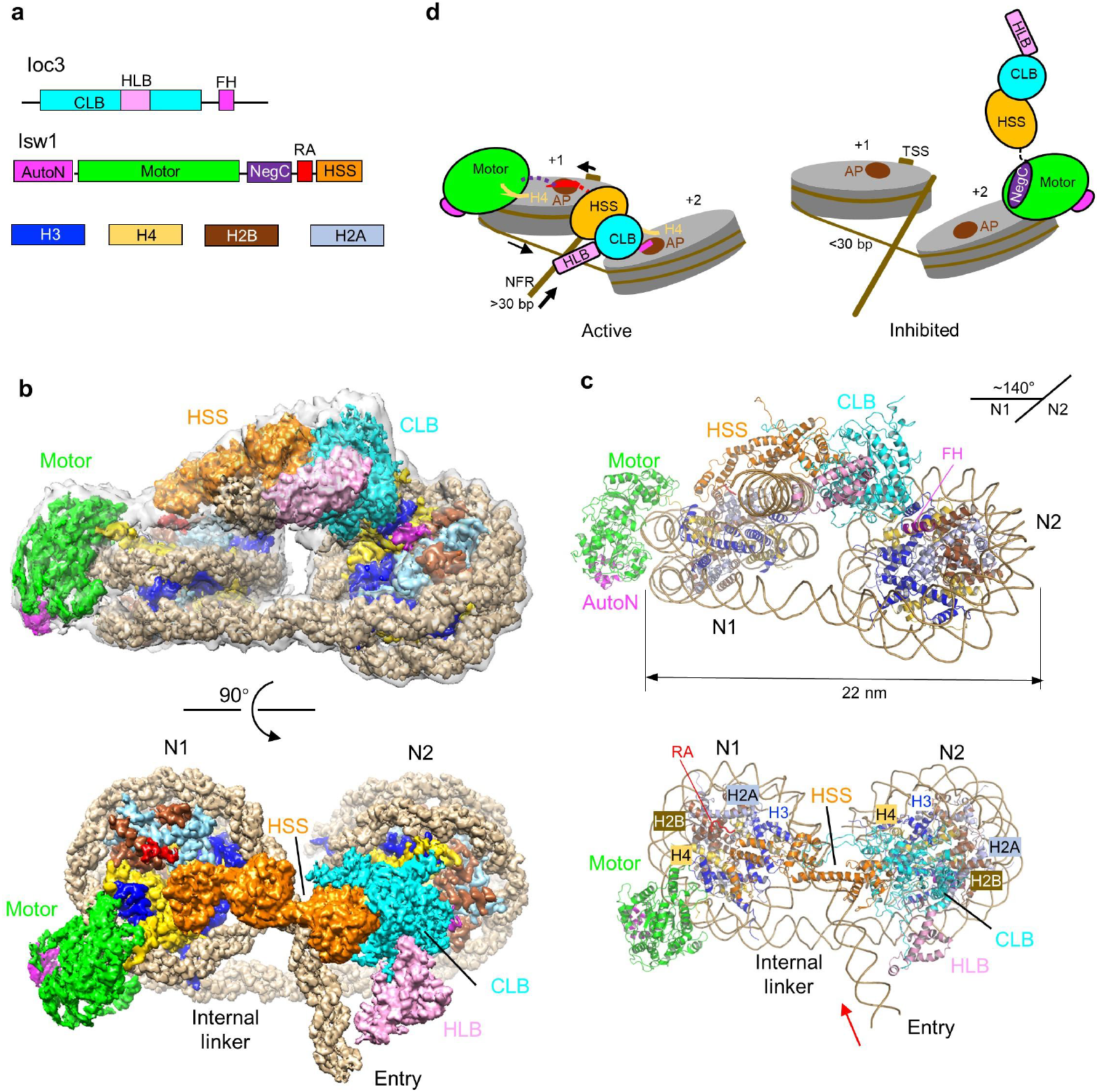
Overall structure of the ISW1a complex bound to the dinucleosome. (**a**) Domain organization of the subunits of the ISW1a complex, and the histones. Proteins are color coded. NegC is not structurally resolved in the active complex. (**b**) Two views of the high-resolution composite maps of the ISW1a-dinucleosome complex. The molecular envelop is shown in the top panel. (**c**) Two views of the ribbon model of the ISW1a complex bound to the dinucleosome. (**d**) Model of ISW1a action on the +1 and +2 nucleosomes of a promoter region. NFR, nucleosome free region; TSS, transcription start site; AP, acidic patch. The arrows indicate the direction of the DNA translocation. The motor binds to, and slides the +1 nucleosome towards the NFR, whereas it is inhibited when bound to the +2 nucleosome with a short linker DNA (<30 bp).

In this study, we report the cryoEM structure of the ISW1a complex bound to the dinucleosome, which illustrates the ISW1a mechanism, and reveals a positive role of the neighboring nucleosome in regulation of chromatin remodeling.

## Overall structure of ISW1a bound to the dinucleosome

Isw1 (residues 69-1129) including all of the functional domains, and Ioc3 (residues 127-749) including the CLB and HLB domains, from *Saccharomyces cerevisiae* were used to form the ISW1a complex, Fig. 1b. The canonical promoter chromatin in yeast cells contains the NFR and downstream nucleosome arrays with an average linker DNA length of ~20 bp^33^. To mimic the architecture of promoter proximal nucleosomes, dinucleosome 30(N1)20(N2)10, which contains internal linker DNA of 20 bp between the two nucleosomes (N1 and N2), and 30 and 10 bp external DNA at either ends, was used as the substrate to capture the action of the motor at the initial engagement state. The ISW1a complex was mixed with the dinucleosome in the presence of the stable ATP analog (ADP-BeFx), and subjected to single-particle cryo-EM analyses (Fig. S1 and Table S1). The structure of the ISW1a complex bound to the dinucleosome was determined at an overall resolution of 5.4 Å, and local maps around N1 (bound with the RA motif), N2 (bound with the finger helix), N1-motor, and N2-Ioc3, were locally refined to 3.2, 2.9, 3.1, and 3.1 Å, respectively (Fig. 1b and Fig.S2).

The N1 and N2 nucleosomes were recognized by the Isw1 motor and Ioc3 subunit, respectively. As reported before^17^, the HSS domain of Isw1 interacts with the CLB domain of Ioc3 to assemble the DNA-binding module, which binds to the external DNA of the N1 nucleosome at the entry side. The dinucleosomes bound by ISW1a are locked into a compact, zigzag conformation, with a width of ~22 nm and a twist angle of ~140°, Fig. 1c. This conformation is more compressed than the structure of the 167-tetranucleosomes^34^, Fig. S3, which have the same internal linker DNA length but a width of 25 nm. Nucleosome fibers can adopt different conformations, and our dinucleosome structure is similar to the reported structures of the dinucleosome with an internal linker DNA length of 29 bp^35^, Fig. S3, suggesting that the dinucleosome- binding mode of ISW1a works with variable internal linker DNA lengths.

## Recognition of the N1 nucleosome by Isw1

Consistent with the previous studies of Isw1 motor fragment^30^, the motor domain of the ISW1a complex bound to the nucleosome at the super helical location 2 (SHL2), Fig. 2a. AutoN was released from the autoinhibited conformation, but remained bound to the motor domain, and the structure of the NegC domain was not clearly detected.

**Figure 2.**
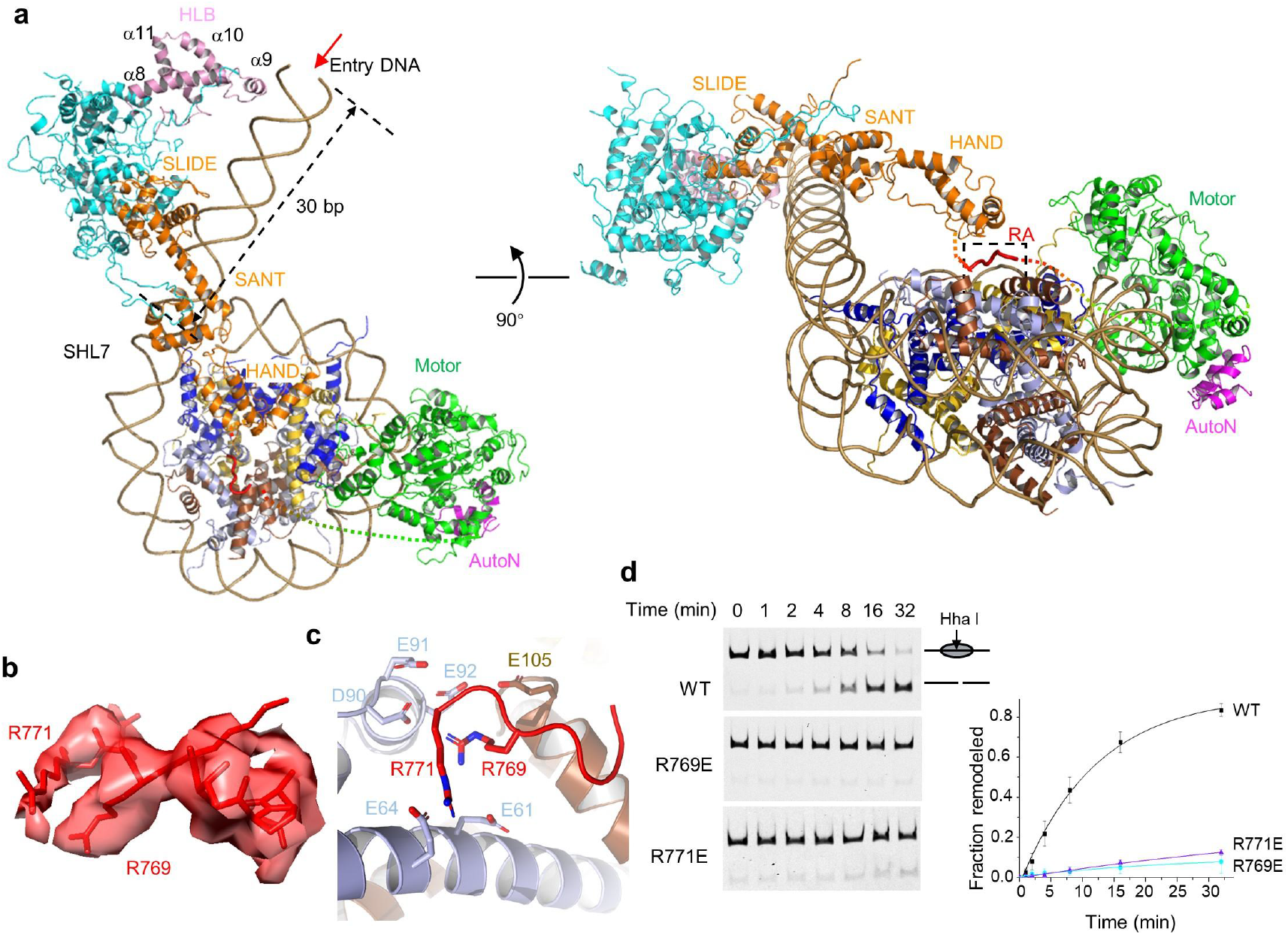
Structure of the ISW1a complex bound to the N1 nucleosome. (**a**) Two views of the structure of the ISW1a complex bound to the N1 nucleosome. The boxed RA region is enlarged for analysis in (c). The arrow indicates the direction of DNA translocation (into the N1 nucleosome). (**b**) Local cryoEM density of the arginine anchor motif of Isw1. (**c**) Recognition of the H2A-H2B acidic patch by two arginine anchors of Isw1. (**d**) Chromatin remodeling activity of the WT and two arginine anchor mutants of the ISW1a complex towards the 100N100 mononucleosome. Quantifications of the remodeling activity are shown on the right. Error bars indicate SD (n=3). The initial reaction rates (mini^-1^) estimated by fitting and data are 0.076, 0.003, 0.003 for WT, R769E, R771E, respectively.

A highly conserved polybasic motif of the ISWI family enzymes recognizes the acidic patch of the nucleosome, which is important for the chromatin remodeling activity^29,36–38^. Despite extensive studies^30–32^, the structural basis of acidic patch recognition by ISWI remains unclear. The 3.2 Å-resolution local map of the N1 nucleosome resolved two arginine residues binding to the acidic patch, Fig. 2b. Based on the structural orientation of the polybasic motif, which connects the N-terminal motor domain at SHL2 to the C-terminal HSS domain at the nucleosome edge, we assigned the two arginine residues as Arg769 and Arg771, Fig. 2c. In particular, Arg769 functions as the canonical RA that interacts with the negatively charged pocket formed by Glu61, Asp90, Glu91 and Glu92 of H2A, and Glu105 of H2B^39^. Consistent with these assignments, relative to the wild type (WT) enzyme, the R769E and R771E mutations reduced the basic nucleosome sliding activity by factors of >20 in a restriction enzyme accessibility assay, in which the 100N100 mononucleosomes with 100 bp DNA at both ends were used as the substrate, Fig 2d. We also tested the activity using a nucleosome spacing assay, in which the closely packed 50N15N50 dinucleosomes were moved to more evenly spaced, fast-migrating products^25^ (Fig. S4a). The RA mutations of Isw1 considerably diminished the activity of the complex, Fig. S4b. The RA residues of Isw1 are highly conserved, suggesting the human homologs recognize the nucleosome through a similar mechanism^29^.

Whereas the motor domain binds at SHL2, the RA motif anchors on the acidic patch. These two binding elements are >70 Å away from the each other (Fig. S5), and put tension on the connecting sequence of the NegC domain. The strain with a long disordered NegC is expected to be much lower than the one associated with a fully folded NegC (Fig. S5c). The structure provides the mechanism to destabilize the folding of the inhibitory NegC domain^27,28^, and thus the activation of the motor.

## Structure basis of ISW1a in nucleosome spacing

The ISW1a complex promotes the formation of nucleosome array with spacing of ~30 bp ^18,22–24^, and the HSS-Ioc3 module was proposed to act as the protein ruler that binds two DNA segments to set the nucleosome spacing^17^. Yet, the binding of the HSS-Ioc3 module to the linker DNA of the nucleosome was not clearly detected before, and its orientation relative to the motor was not defined.

In the cryoEM structure of the ISW1a complex, the secondary structural elements of the HSS-Ioc3 module were clearly resolved, which adopts a conformation similar to the crystal structure^17^, Figs. S2e and S6. Different from the previous model, the HSS-Ioc3 module binds to the mobile nucleosome (defined by the engagement of the motor) at the same DNA gyre as the motor domain, and interacts with the entry DNA, without contacting the second DNA segment from the neighboring nucleosome, Fig. 2a. In contrast to the bent DNA structures detected before^17^, the entry DNA bound by the HSS-Ioc3 module adopts a straight conformation, Fig. S6a. The HAND domain of Isw1 does not interact with the DNA, and is poised above the disk face of the nucleosome, close to the RA motif. The SANT domain interacts with the first minor groove of the linker DNA appearing at the nucleosome edge, and the SLIDE domain binds ~10 bp upstream. The α9 of the HLB domain of Ioc3 binds, through its polybasic surface, to the major groove of the entry DNA located ~20 bp further upstream, Fig. S6b. Therefore, the combined actions of HSS and Ioc3 measure the entry DNA length of ~30 bp from the nucleosome edge. Our findings are consistent with the previous studies showing that the HLB domain is important for nucleosome organization in cells^26^.

The RA motif and the HSS-Ioc3 module probably bind the nucleosome in a cooperative manner, as suggested by the lack of stable binding of the RA motif in the previous motor structure^30^. These two auxiliary elements stabilize the motor domain binding at the same face of the mobile nucleosome. The Isw1 motor translocates the entry DNA into the nucleosome (Fig. 2a), until the entry linker DNA is too short for the stable HSS-Ioc3 binding. This model provides the mechanism of ISW1a in nucleosome spacing, which is in line with the idea of protein ruler, but without the requirement for the binding of the second DNA fragment by the CLB domain as suggested before^17^.

## Recognition of the N2 nucleosome by Ioc3

Unexpectedly, we found that Ioc3 binds to the disk face of the N2 nucleosome, Fig. 3a. Docking the crystal structure of Ioc3 into the cryoEM density map revealed that a positively charged surface comprising Lys292, Lys293, Arg373 and Lys377 of the CLB domain is close to the minor groove DNA at the SHL1.5 of the N2 nucleosome^17^, Fig. 3b. Furthermore, the high-resolution local map clearly resolved the adjacent polybasic N-tail of H4, which binds to the negatively charged surface of Ioc3, Fig 3c. Specifically, Arg23 of H4 interacts with Gln251 and Asp252 of Ioc3, and Arg19 of H4 is close to Asp240 of Ioc3. The conformation of H4 is stabilized further by the interactions of Lys16, Arg17 and Lys20 with the DNA at SHL2.5. Notably, the C-terminal helix of Ioc3 (referred as to the finger helix) binds to the H2A-H2B acidic patch of the N2 nucleosome, Fig 3d. Four arginine residues, Arg735, Arg738, Arg739 and Arg742 cluster on one side of the finger helix, with similar orientations and compositions to those of the finger helices of the Snf5/Sfh1/SMARCB1 subunits of the SWI/SNF family complexes^40–42^, supporting functional convergence on a common mechanism of nucleosome recognition, Fig. S7.

**Figure 3.**
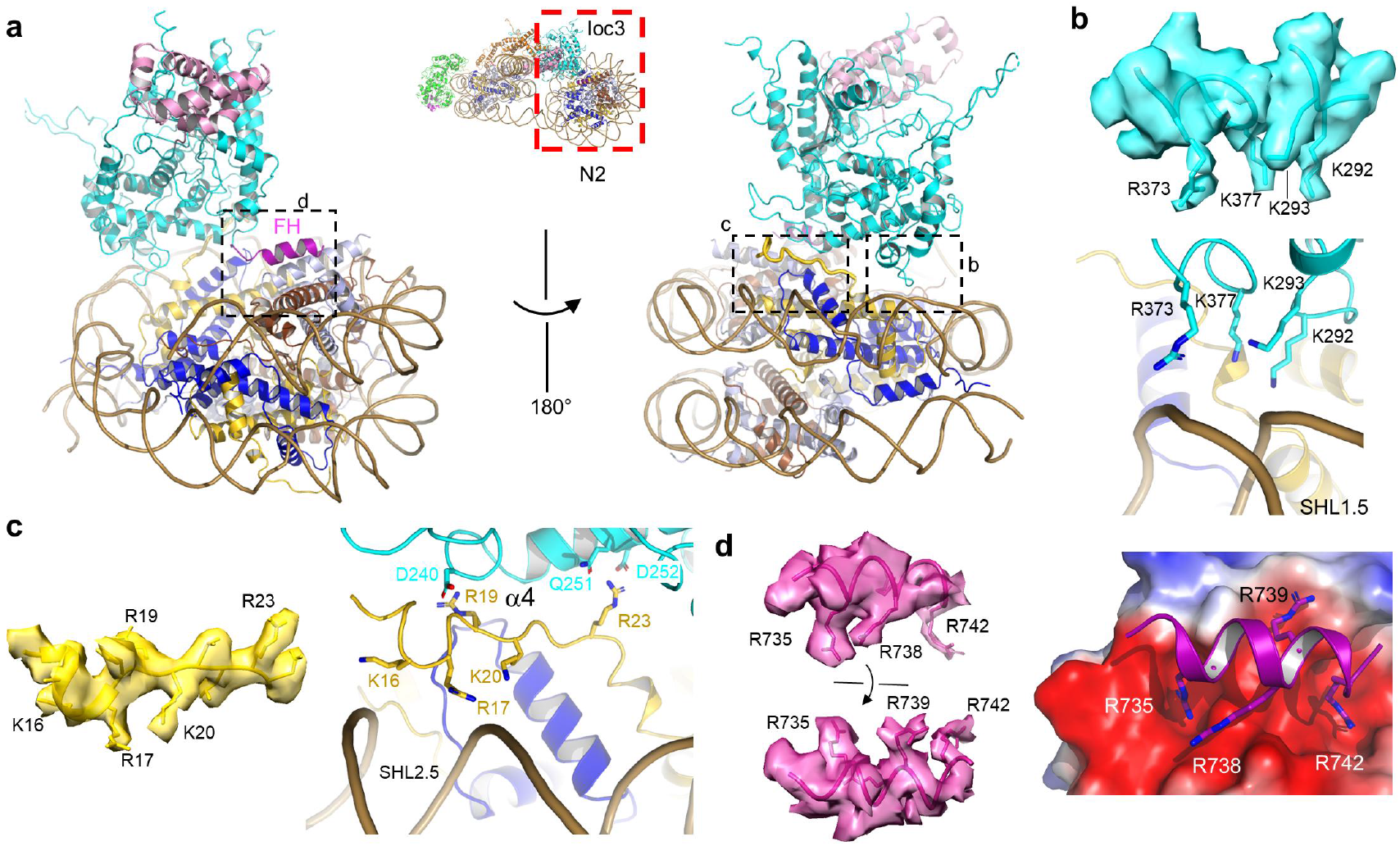
Structure of Ioc3 bound to the N2 nucleosome. (**a**) Two views of the structure of Ioc3 bound to the N2 nucleosome. The left panel is oriented similar to the inset. The black-box regions are enlarged for analysis in (b-d). (**b**) Interaction between Ioc3 and the nucleosomal DNA at SHL1.5. The local EM density map of Ioc3 is shown in the top panel. (**c**) Recognition of the H4 tail by Ioc3. The local EM density map of the H4 tail is shown on the left. (**d**) Recognition of the H2A-H2B acidic patch of the nucleosome by the finger helix of Ioc3. The local EM density of the finger helix is shown on the left. Electrostatics was calculated with Pymol.

## Nucleosome recognition by Ioc3 is important for nucleosome spacing in vitro

Nucleosome recognition by Ioc3 is intriguing. To determine its significance, we first tested the nucleosome sliding activity toward the 100N100 mononucleosomes. The Ioc3 mutants Q251K/D252K (Ioc3-H4), K292E/K293E (Ioc3-DNA), and R739E/R742E (Ioc3-AP) were constructed to disrupt the interactions with the H4 tail, the nucleosomal DNA, and the H2A-H2B acidic patch, respectively. Consistent with its role in binding to the neighboring nucleosome but not the mobile nucleosome, these Ioc3 mutants showed activities comparable to the WT enzyme toward the 100N100 mononucleosome, Fig. 4a. Similar observations were found using a mononucleosome centering assay, Fig. S4c. The data indicate that recognition of the N2 nucleosome by Ioc3 is not important for remodeling the substrate with just one nucleosome.

**Figure 4.**
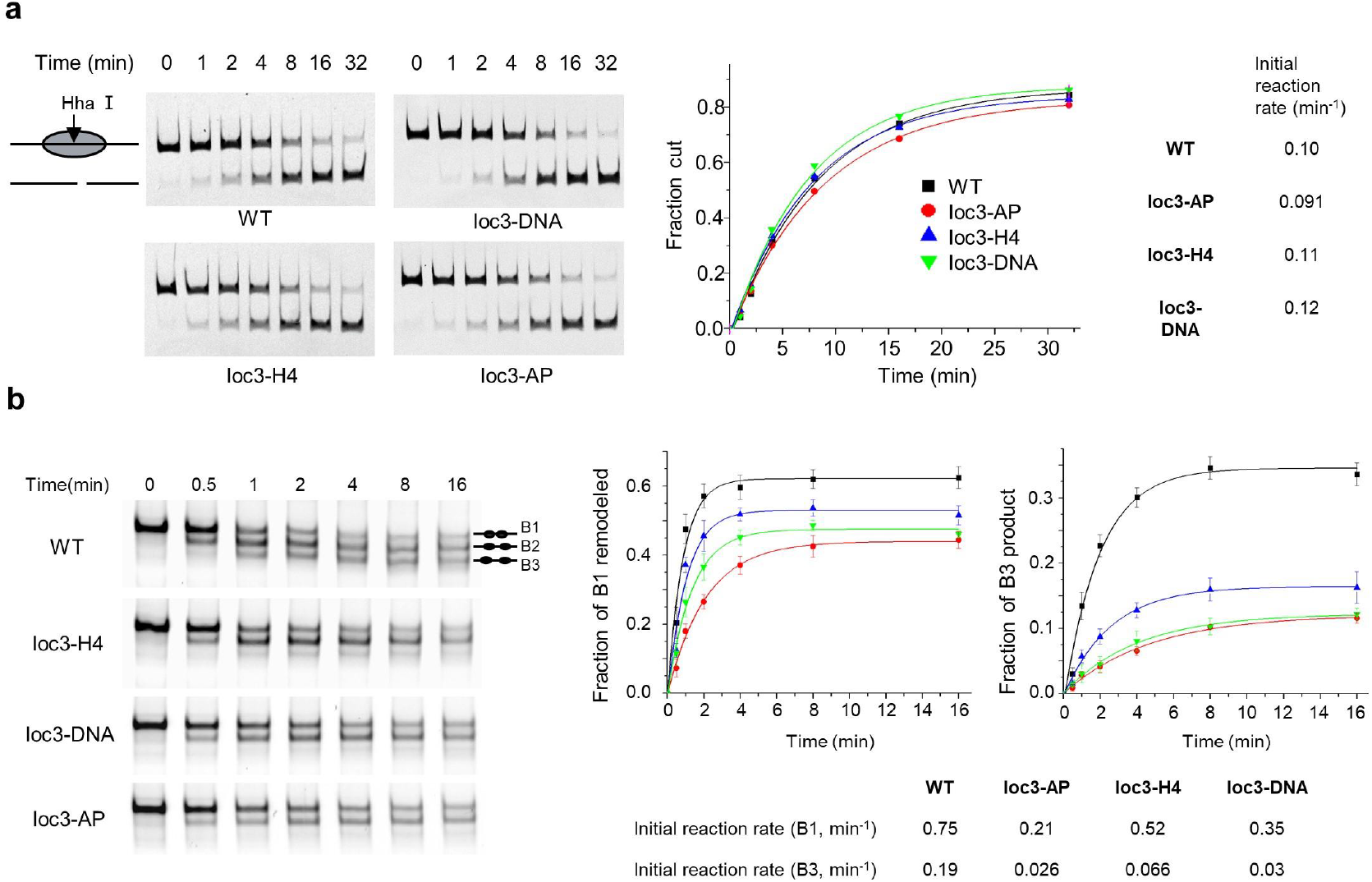
Nucleosome recognition by Ioc3 is important for nucleosome spacing in vitro. **(a)** Chromatin remodeling activity of the WT and three Ioc3 mutants of the ISW1a complex toward the 100N100 mononucleosome. Quantifications of the remodeling activity are shown on the right. Error bars indicate SD (n=3). The initial reaction rates are estimated by fitting and data, and shown on the right. **(b)** Chromatin remodeling activity of the WT and three Ioc3 mutants of the ISW1a complex towards the 45N15N45 dinucleosome. Quantifications of the remodeled substrate (B1), and the formation of final product (B3) are shown on the middle and right panels, respectively. Error bars indicate SD (n=3). The initial reaction rates are estimated by fitting and data, and shown at the bottom.

Using dinucleosome 45N15N45 as the substrate, the Ioc3 mutants displayed notable defects, Fig. 4b. The ISW1a complex remodeled the dinucleosome (band B1) into two main species (bands B2 and B3). The intermediate product (band B2) formed first, and the more evenly spaced nucleosomes (band B3) appeared later. Comparison of the initial rates of the substrate (B1) remodeled indicated that relative to the WT complex, the three Ioc3 mutants showed lower activities (Fig. 4b). Comparison of the formation of the product of evenly spaced nucleosomes (B3) indicated even more noticeable reductions of the activity. The data support that recognition of the neighboring nucleosome is important for the maximal spacing activity. The losses of the spacing activity of the Ioc3 mutants were also observed using a different dinucleosome substrate (10N10N60), Fig. S4d. Together, the data suggest that nucleosome recognition by Ioc3 is not essential for the basic translocation activity to slide the mononucleosome, but that it is important for the spacing activity to sense the neighboring nucleosomes in the context of chromatin environment.

## Nucleosome recognition by Ioc3 is important for cell fitness and chromatin organization in vivo

We then tested the function of nucleosome recognition by Ioc3 inside the cells. Compared with the WT cells, the Ioc3 deletion (ΔIoc3), Ioc3-AP and Ioc3-H4 mutant cells showed similar growth under normal conditions. Fig. 5a. However, under the stressed conditions, in the presence of the genotoxic agent methyl methanesulfonate (MMS) at 37 °C in particular, the mutant cells displayed slow growth phenotype, suggesting defects in the expression of stress response genes.

**Figure 5.**
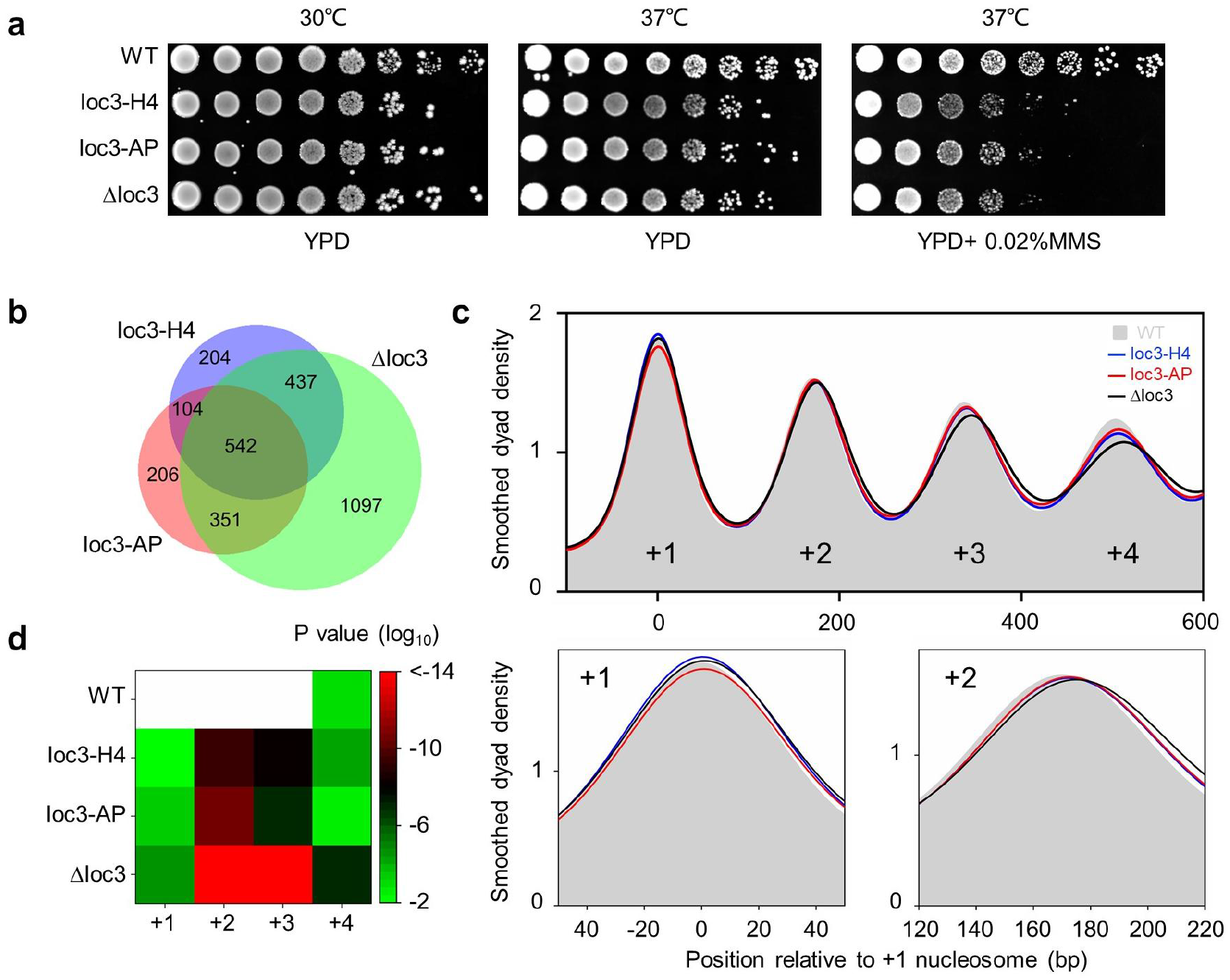
Recognition of the N2 nucleosome by Ioc3 is important for the functions of the ISW1a complex in cells. (**a**) Growth assays of the WT and Ioc3 mutants yeast cells under different conditions. Three independent assays were performed, and a representative result is shown. (**b**) Venn diagram showing the overlap of the genes with at least one shifted nucleosome in the +1 to +4 positions (p < 0.01). The 542 overlapping genes were chosen for further analysis. (**c**) Promoter nucleosome positioning of the WT and Ioc3 mutant yeast cells. The positioning of the promoter proximal nucleosomes of 542 genes commonly altered in the Ioc3 mutant cells are shown. The regions around the +1 and +2 nucleosomes are enlarged for a clearer view of the nucleosome shifts. (**d**) Statistical significance of the nucleosome shifts in the mutants shown by the log10 p-values (Wilcoxon-Mann-Whitney test) as a heat-map. The positions in WT cells were used as the reference. The shifts of an independent WT sample are used as the control.

Finally, we characterized the genomic nucleosome organization of the cells using MNase-seq, Figs. 5b–5d, S8 and S9. Consistent with previous studies^26^, the WT cells displayed well positioned promoter nucleosomes, and deletion of Ioc3 generally shifted the promoter nucleosomes away from the transcription start sites, with the +1 and +2 nucleosomes shifting 2 and 4 bp, respectively, Figs. 5c, S8a and S8b. The Ioc3-AP and Ioc3-H4 mutants elicited the same trend in nucleosome movement as observed in the ΔIoc3 mutant, with the +1 and +2 nucleosomes of the 542 commonly affected genes shifting downstream by an average of 1 and 2 bp, respectively (Fig. 5b–5c, with the statistical significance shown in Fig. 5d). The point mutants induced smaller effects than deletion of the whole Ioc3 gene, suggesting that multiple elements of Ioc3 work together in nucleosome recognition. These results were exemplified at several loci, Fig. S9. Together, the data indicate that sensing the neighboring nucleosomes by Ioc3 is important for chromatin organization in cells.

## Discussion

In this study, we determined the structure of ISW1a bound to the dinucleosome, which not only provides an integrated picture of the ISW1a mechanism, but also adds a new dimension to the regulation of nucleosome remodeling in the context of higher-order chromatin structure.

The previous studies illustrate DNA recognition by the isolated DNA-binding module and nucleosome recognition by the motor domain^17,30^. The current structure reveals an integrated picture how the ISW1a complex function as a unit to interact with the nucleosome substrate through multiple elements to set up nucleosome spacing, Fig. 1d. The motor domain binds to the mobile nucleosome (N1). The RA motif anchors on the surface of the acidic patch of the mobile nucleosome, which probably imposes strains on the preceding NegC domain to unfold. The following HSS domain binds to the entry linker DNA by ~30 bp close to the nucleosome edge. Thus, the motor, the RA motif, and HSS-Ioc3 module collectively bind to the same face of the mobile nucleosome, which provides the structural basis of linker DNA sensing of the enzyme. Remarkably, the Ioc3 subunit recognizes the neighboring N2 nucleosome through interactions with the acidic patch, the H4 tail and the nucleosomal DNA, which enhances the spacing activity of the complex.

In contrast, engagement with the nucleosome with a short length DNA (<30 bp), Fig. 1d right panel, destabilizes the binding of the HSS-Ioc3 module, which results in folding back of NegC and thus inhibition of the motor, as suggested before^27,28^. The position of the HSS-Ioc3 module in the inhibited state is not clear. Therefore, a long linker DNA >30 bp promotes the ISW1a activity, whereas the motor is inhibited by the folded-back NegC as the linker DNA gets shorter.

In this model, the new regulatory mechanism of Ioc3 does not impose directly on the motor, and thus does not alter the fundamental sliding activity of the Isw1a complex to slide the mononucleosome. Instead, it interacts with neighborhood nucleosome, and specially changes the spacing activity towards nucleosome arrays. In contrast, the RA motif promotes the basic nucleosome sliding activity of the motor.

This mechanism of ISW1a is line with the previous idea of “protein ruler” ^17^, but differs in several aspects. The new model includes a novel role of Ioc3 in sensing the nearby nucleosome, whereas it does not involve the binding to the secondary I/E DNA strand as proposed before^17^. The new model also includes a breaking mechanism of NegC inhibition to actively stop the remodeling reaction as the linker DNA gets too short. Therefore, the nucleosome arrays along the genomic DNA not only confine the linker DNA lengths ^8,17^ but also provide a positive signal to regulate the remodeling activity. Recognition of the mobile nucleosome by the motors is well known^24,27–29,36,39,40,43^. The ISW1a complex is the first chromatin remodeler that recognizes the neighboring nucleosome ^25,26^. Our work provides mechanistic insights into the regulation within a chromatin environment beyond the individual nucleosome.

The positions and spacing of the promoter nucleosomes are controlled by the intrinsic property of the DNA sequence, and the collective actions of DNA-binding factors and chromatin remodelers^6,16^. In particular, the actions of ISW1a and RSC antagonize each other, in which ISW1a slides the promoter nucleosomes to close the NFR, whereas RSC opens up the NFR^13,15,16,20^. Our model provides insights into their mechanism of action, Fig. 1d. The transcription start sites of the genes in yeast generally reside ~10 bp inside the +1 nucleosome^2^. The 20 bp inter-nucleosome linker DNA length is too short for the HSS-Ioc3 module of ISW1a, which instead binds to the NFR DNA at the edge of+1 nucleosome, allowing the Isw1 motor to land at SHL2 of the same DNA gyre. Recognizes the disk face of the adjacent +2 nucleosome by Ioc3 further promotes the activity. Therefore, ISW1a senses the NFR, the +1 nucleosome, the inter-nucleosome linker DNA, and the +2 nucleosome in remodeling the promoter nucleosomes. The firing of the Isw1 motor translocates the NFR DNA into the +1 nucleosome (equivalent to slide the +1 nucleosome towards the NFR), further occludes the transcription start site, and potentially represses the expression of a gene. This is in contrast with the RSC complex, in which the auxiliary subunits Rsc3/30 recognize the NFR DNA, but the motor lands at the distal DNA gyre^40,43^. RSC slides the nucleosome away from the NFR and opens up the promoter^15,44,45^. Thus, both ISW1a and RSC bind the NFR DNA, but the specific architectures of the remodeling complexes place the motor domains to different positions of the +1 nucleosome, and function in closing and opening up the promoter chromatin, respectively.

## METHODS

### Protein expression and purification

We cloned the genes of Isw1 (residues 69–1,129) and Ioc3 (residues 127–749) from *S. cerevisiae* genomic DNA, then inserted them into the pRSF Duet vector containing a SUMO tag at the N terminus of Ioc3. The point mutants were generated by Quikchange mutagenesis. All constructs were confirmed by DNA sequencing.

The ISW1a complex was overexpressed in the *Escherichia coli* expression strain Rosetta (DE3) induced with 0.5 mM isopropyl-beta-D-thiogalactoside at OD600~0.8, and cultured at 18 °C overnight. Cells were harvested by centrifugation at 4,000 r.p.m. (Beckman, Rotor JA4.2) for 20 min, resuspended in 50 mM Tris-HCl, 300 mM NaCl, 5% Glycerol, 10 mM imidazole and 1mM phenylmethanesulfonyl fluoride, pH 8.0 and then disrupted by a high-pressure homogenizer (ATS). After high-speed centrifugation, the supernatant was loaded on a Ni–NTA column and allowed to flow through by gravity. After a washing step with 30 mM imidazole, the complex was eluted with 20 mM Tris-HCl, 300 mM NaCl, 200 mM imidazole, and pH 8.0. The His6-SUMO tag was cleaved by SUMO Protease (Ulp) at 4 °C for 3 h. The complex was further purified on an ion-exchange column (Source-15Q, GE Healthcare) with Tris-HCl buffer, pH 8.0, and then subjected to gel-filtration chromatography (Superdex-200, GE Healthcare) in buffer containing 10 mM HEPES, 300 mM NaCl, 20% Glycerol, 5 mM DTT and pH 7.5. The purified complex was concentrated to 10 mg/ml and stored at −80 °C. The mutants of ISW1a complex were purified similarly, and concentrated to 3-6 mg/ml.

### Nucleosome reconstitution

Nucleosome core particles (NCP) were reconstituted with Xenopus histones and the “601” DNA as described previously^46^. To assemble the mononucleosomes, the ‘601’ DNA fragments with 80 bp linker DNA at one end (0N80), and with 100 bp linker DNA at both ends (100N100) were used. To assemble the dinucleosomes with different internucleosome spacing, two copies of “601” sequences were constructed into pUC57 plasmid with 15 bp (50N15N50) and 20 bp (30(N1)20(N2)10) linker sequences.

### Nucleosome remodeling assay

Nucleosome centering assays were performed basically as described with 20 nM Cy5-labeled mononucleosomes, and 80 nM ISW1a complex^28^. To remodel the 0N80 mononucleosome, reactions were performed in buffer (50 mM KCl, 20 mM HEPES, 5 mM MgCl_2_, 0.1 mg/ml bovine serum albumin, 5 % glycerol, 1 mM DTT, and pH 7.5) with 10 μM ATP at 25 °C, and stopped with 800 ng/μl sperm DNA at the indicated time points. The products were resolved with 8% native acrylamide gels, 0.25×TBE at 4°C and 150V for 150 min.

To remodel the dinucleosomes (45N15N45, 50N15N50, and 10N10N60) in the spacing assays, reactions were performed in buffer (100 mM KCl, 20mM HEPES, 5 mM MgCl_2_, 0.1 mg/ml bovine serum albumin, 5% glycerol, 1 mMDTT, and pH 7.5) with 10 μM ATP and 400 μM ATP-γ-S at 25 °C, and stopped with 800 ng/μl sperm DNA. The products were analyzed on native 5.5% polyacrylamide gels, 0.25×TBE at 4°C and 120V for 180min.

The REAA assays were performed as described^28^. Cy5-labelled 100N100 NCP (5 nM) and 10 nM of ISW1a WT and various mutant complexes were incubated at 25 °C with 3 mM ATP and 100 U of Hha I in the reaction buffer (50 mM KCl, 20 mM HEPES, 5 mM MgCl_2_, 0.1 mg/ml bovine serum albumin, 5% glycerol, 1 mM DTT, and pH 7.5). Fractions were taken at various time points and quenched with 2× stop buffer (20 mM Tris, 0.6% sodium dodecyl sulphate (SDS), 40 mM EDTA, pH 8.0 and 0.1mg/ml proteinase K). The reaction mixtures were incubated at 55 °C for 20 min to deproteinate the samples. Fractions were running on 8% polyacrylamide gels for 100 min at 100 V.

The Cy5-labeled DNA bands were detected using a Typhoon9410 imager (GE Healthcare). Band intensities were quantified in Quantity One software. The fraction of nucleosome/DNA remodeled at a given time point was determined by the ratio of remodeled nucleosome/DNA to total nucleosome/DNA intensity in the same lane of the gels. Considering the final products often reached to different levels, the initial reaction rates were fit to a single exponential decay using Origin9.2, and used to compare the remodeling activities.

### Electron microscopy sample preparation and Data Collection

The ISW1a-NCP complexes were obtained by mixing 6 μM protein with 4 μM dinucleosome 30(N1)20(N2)10 in the presence of ADP-BeFx. The complex was dialyzed to buffer containing 50 mM NaCl, 10 mM Hepes, 1 mM MgCl_2_, 0.25 mM ADP, 0.5 mM BeSO_4_, 4 mM sodium fluoride and pH 7.5. The resultant mixture was then stabilized and purified using the GraFix method as described^47^.

Negative staining of the ISW1a-nucleosome complex was performed with 2% uranylacetate. Grids were examined using an FEI microscope operated at 120 kV, and the images were recorded using a charge-coupled device (CCD) camera (Gatan). For cryo-EM grids preparation, Quantifoil gold R2/1 grids with 200 mesh size were subjected to glow discharge in air for 15 s using a PDC-32G-2 Plasma Cleaner set to a low power. A drop of 4 μl sample was blotted for 3.5 s at −2 force before being plunge-frozen in liquid ethane with a FEI Vitrobot IV at 8 °C and 100% humidity. Grids were screened using Talos Arctica operated at 200 kV. Cryo-EM data were collected using a Krios G3i operated at 300 kV equipped with a K3 direct electron detector and GIF Quantum energy filter (Gatan), at a magnification of 81,000X for a final pixel size of 0.5412 Å/pixel with the defocus values ranging from −1.4 to −1.8 μm. The total electron dose was 50 e-/Å2 fractionated in 32 frames (exposure time 2.56s).

### Image processing

The initial 3D model of the ISW1a-nucleosome complex was constructed using Relion3.0^48^. The ISW1a-nucleosome datasets were collected from two sessions. All the 16,488 dose-fractionated image stacks were aligned using MotionCor2^49^ with twofold binning, and the CTF parameters were estimated using CTFFIND4^50^. Particle picking, two-dimensional (2D) classification and three-dimensional (3D) classification were carried out in Relion3.0. The initially picked particles were extracted with fourfold binning to increase signal to noise ratios. After multiple rounds of 2D classification, a total of 3,070,142 particles were selected from the dataset1 and 2,699,804 particles were selected from the dataset2. After seed-facilitated 3D classification^51^, 1,058,952 raw particles were selected and 311,464 particles with clear N2-Ioc3 features were picked for further focused 3D classification. The best 212,066 particles were refined to 3.1 Å for the N2-Ioc3 module. To remove broken particles, several rounds of 3D classification were applied and the selected 292,854 particles were further classified to remove particles without the intact Ioc3. The re-extracted 208,137 particles without binning were refined with soft masks of N2-Ioc3 and N1-motor, respectively. After multiple rounds of focused 3D classification, 166,775 particles were selected for the N2 bound to the Finger Helix of Ioc3 (N2-FH) and refined to 2.9 Å. Similar pre-processing was performed on the dataset2, and 219,554 particles were selected. To improve the N1-motor module, the 170,126 particles from the two datasets resulted from multi-round focused 3D classification were combined. N1 bound to the arginine anchor of Isw1 (N1-RA) was refined to 3.2 Å, and N1 bound to the Isw1 motor was further improved to 3.1 Å after masked 3D classification. The estimated overall resolution of ISW1a-nucleosome complex was 5.4 Å after re-extraction, 3D auto-refinement and post-processing. To clarify the interaction between Ioc3 and DNA, the signals outside the DNA-Ioc3-HSS density were subtracted and focused 3D classification resulted in 66,852 particles for global refinement. The particles with more accurate angular alignment were performed signal subtraction on the DNA-Ioc3-HSS density again and the selected 41,577 particles were refined to 6.5 Å with clear secondary structures.

### Model building

The initial model was built by fitting the maps in Chimera using the known structures, including ISWI-nucleosome complex (PDB code 6JYL)^30^, the crystal structure of Ioc3-HSS (PDB code 2Y9Z)^17^ as the templates. The atomic models for the rest of the molecules were built manually in Coot. The structures were refined using Phenix with secondary structure constrains^52^.

### Yeast genetics

The yeast stain BY4741 (MATa *leu2 ura3 his3 met15 can1*) was used, transformed with lithium acetate method. To generate the Ioc3 mutant strains (Q251K/ D252K, R739E/R742E), the WT Ioc3 gene was first replaced by a DNA fragment carried a URA3 gene, which enabled the strain to acquire the ability to grow on SC/-Ura sodium medium. The URA3 gene was then replaced by the Ioc3 mutants (Q251K/ D252K and R739E/R742E, respectively), and the cells were selected using 5-FOA sodium medium (SC+5-FOA). All the mutants were confirmed by PCR, and DNA-sequencing.

To determine the effect of the mutations on yeast growth, we conducted spot assays. All strains were cultured in YPD for 48 hours to the saturation stage, diluted in YPD to the optical density OD600 of 0.1, and grew to 1 at 30°C. Twenty-fold serial dilutions of the transformants were spotted on the control and drug plates (SC+0.02% Methyl methanesulfonate, MMS) and incubated at 30 °C for 2 days for drug sensitivity testing. All assays were performed in triplicate.

### MNase-seq

Yeast strains were grown to mid-log phase in YPD medium. Mononucleosomal DNA were digested and purified as described except that the samples were not crosslinked by formaldehyde and the digested samples were collected without gel-purification^53^. DNA libraries were prepared using NEBNext Ultra II DNA Library Prep Kit (NEB #E7645S), and sequenced by NovaSeq 6000 System.

Pair-end sequencing reads were trimmed by Trimmomatic and were mapped to sacCer3 reference genome using Bowtie2^54,55^. Reads in the range of 120-180bp were selected, and those mapped to repetitive rRNA locus (chrXII: 451275–469084) were excluded^56^. SAMtools was used to remove possible PCR duplicates^57^. At this step, processed data from two biological replicates were merged, and an equal number of reads (10,345,423 reads) was randomly selected for each sample. Nucleosome positions were identified using algorithm as described for GeneTrack^58^. Briefly. nucleosome midpoint coordinates were smoothed by Gaussian filtering with a standard deviation of 20 bp. The peak of the smoothed curve defines the nucleosome position^59^. If multiple peaks were within ±73 bp, only the highest peak will be retained. A total of ~4,750 genes larger than 560 bp and with at least four well-defined nucleosome positions (interval <250 bp) were considered for downstream analysis to reduce background noise. +1 nucleosome was defined as the nucleosome that was closest to and within ±150 bp of TSS. For nucleosome shift analysis, the Welsh’s t test was performed by comparing the distribution of all reads that map to within ±73 bp of the dyad position of the respective nucleosomes of the mutants and wild-type^20^. When p < 0.01, nucleosomes were considered to have significant shift. 542 overlapping genes with at least one significantly shifted nucleosome were used for calculating median shift. The statistical significance of the median shifts was determined using the Wilcoxon-Mann-Whitney test.

### Intramolecular tension calculated with the worm-like-chain (WLC) model

The force (*F*) required to extend a polymer calculated with the WLC model is given by:

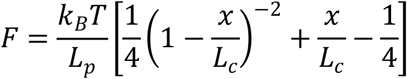

Here, *k_B_T* is Boltzmann’s constant times the absolute temperature; *L_p_* is the persistent length; *L_c_* is the contour length and *x* is the end-to-end distance. The *L_c_* of the NegC is equal to the total number of the disordered residues between the bound motor and the arginine anchor motif multiplied by the distance per amino acid (0.364 nm) as described^60^. The range of *L_p_*=0.5–1.5 nm was used for the calculation, as most of the remodeling assays for ISW1a were carried out under mild ionic strength (20 mM HEPES, 100 mMKCl)^61^.

