## Supplementary Material for "Structure of the ISW1a complex bound to the dinucleosome"

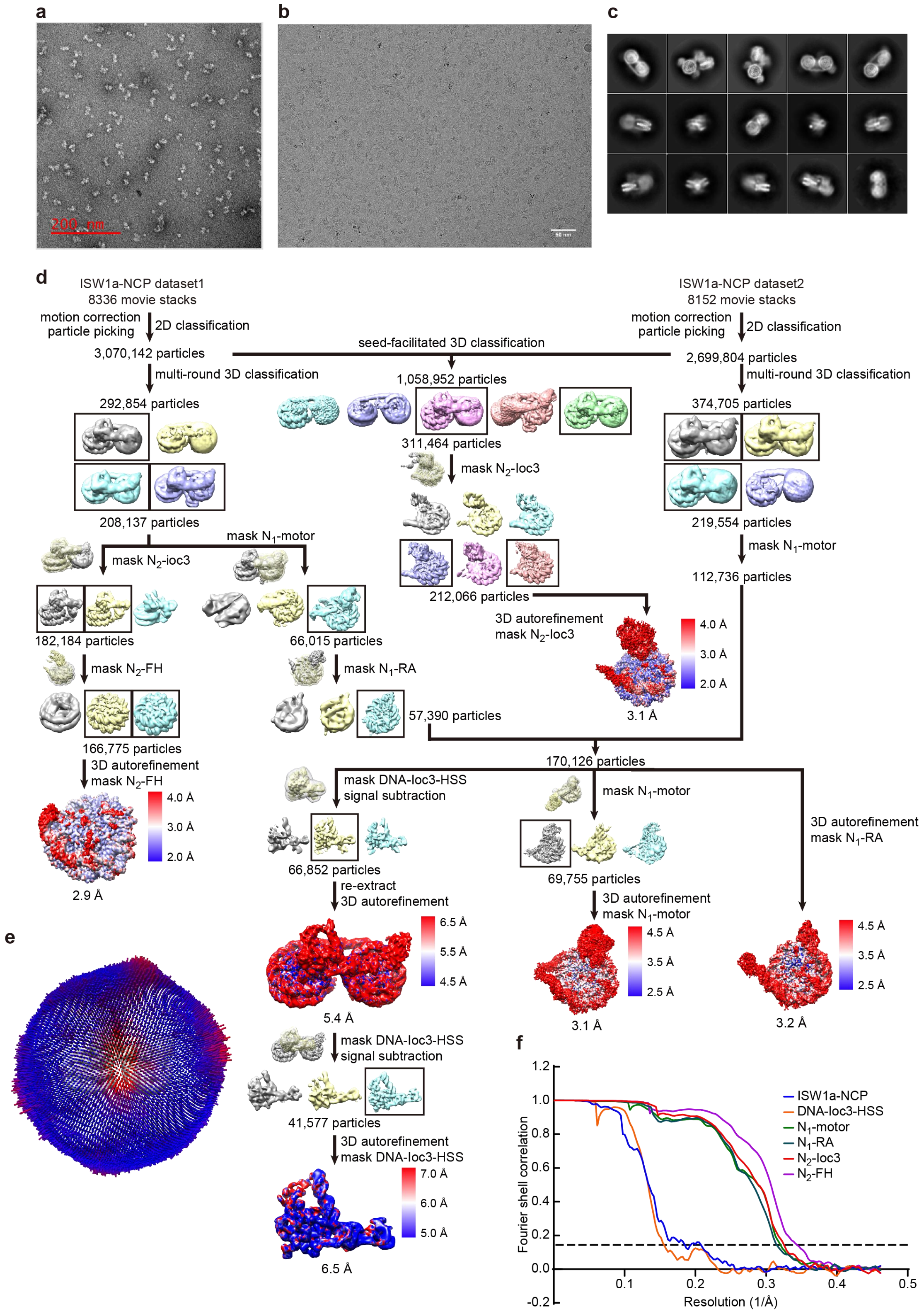
a
b
c
d
ISW1a-NCP dataset1
8336 movie stacks
motion correction
particle picking
2D classification
3,070,142 particles
multi-round 3D classification
292,854 particles
208,137 particles
mask N1-motor
mask N2-ioc3
182,184 particles
66,015 particles
mask N2-FH
mask N1-RA
166,775 particles
57,390 particles
3D autorefinement
mask N2-FH
2.9 Å
ISW1a-NCP dataset2
8152 movie stacks
2,699,804 particles
374,705 particles
219,554 particles
112,736 particles
seed-facilitated 3D classification
1,058,952 particles
311,464 particles
mask N2-ioc3
212,066 particles
3D autorefinement
mask N2-ioc3
3.1 Å
170,126 particles
mask DNA-ioc3-HSS
signal subtraction
66,852 particles
re-extract
3D autorefinement
5.4 Å
41,577 particles
3D autorefinement
mask DNA-ioc3-HSS
6.5 Å
mask N1-motor
69,755 particles
3D autorefinement
mask N1-motor
3.1 Å
3D autorefinement
mask N1-RA
3.2 Å
e
f
Fourier shell correlation
Resolution (1/Å)
ISW1a-NCP
DNA-ioc3-HSS
N1-motor
N1-RA
N2-ioc3
N2-FH

**Supplementary Figure 1 | CryoEM analysis of the Isw1a-dinucleosome complex**. (**a**) Representative negative stain images, (**b**) cryoEM images, and (**c**) 2D classification of the ISW1a complex bound to the dinucleosome. (**d**) Flowchart of the cryo-EM data processing. (**e**) Angular distributions of cryo-EM particles in the final round of refinement of the masked dataset. (**f**) Gold standard Fourier shell correlation (FSC) curves, showing the resolutions of 5.4 Å, 3.2 Å, 2.9 Å, 3.1 Å and 3.1 Å for the ISW1a-dinucleosome complex, N1 (bound with arginine anchors, RAs), N2 (bound with the Finger helix, FH), N1-motor, and N2-Ioc3, respectively.

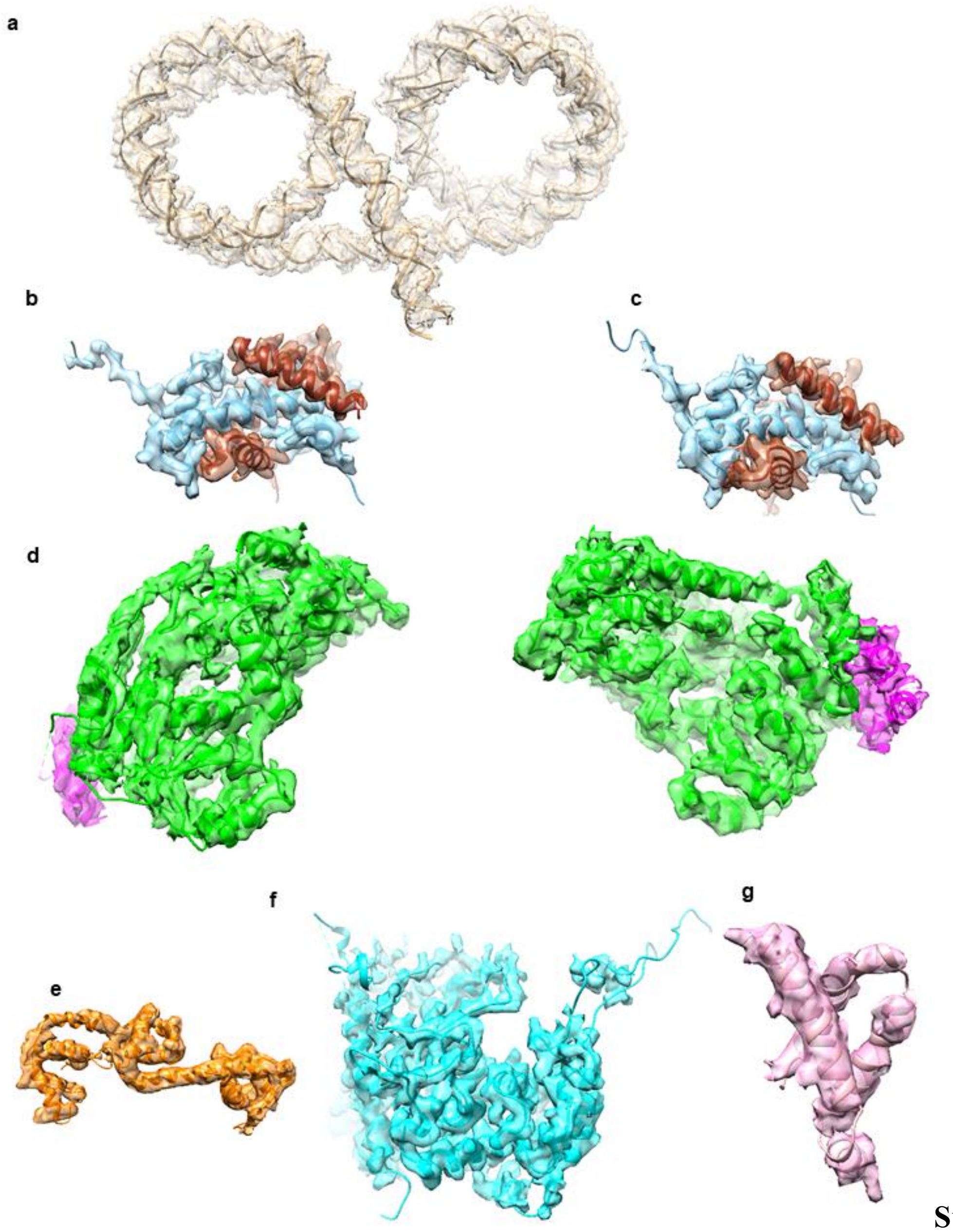


**Supplementary Figure 2 | Local density maps of the Isw1a-dinucleosome complex around** (**a**) nucleosomal DNA; (**b**) H2A-H2B of the N1 nucleosome; (**c**) H2A-H2B of the N2 nucleosome; (**d**) motor domain of Isw1; (**e**) HSS domain of Isw1 (**f**) the CLB domain of Ioc3 and (**g**) HLB domain of Ioc3.

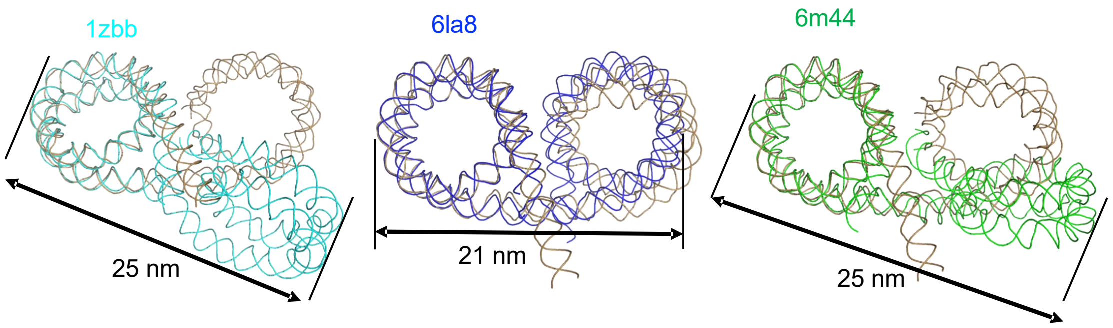


**Supplementary Figure 3 | Comparison of the structures of the dinucleosomes.** The structures of the histone cores are omitted for clarity, and the first histone cores are aligned. The structure determined in this study is in light brown, and PBD codes 1zbb (20 bp linker DNA)[1], 6la8 (29 bp linker DNA)[2] and 6m44 (31 bp linker DNA)[2] in cyan, blue and green, respectively.

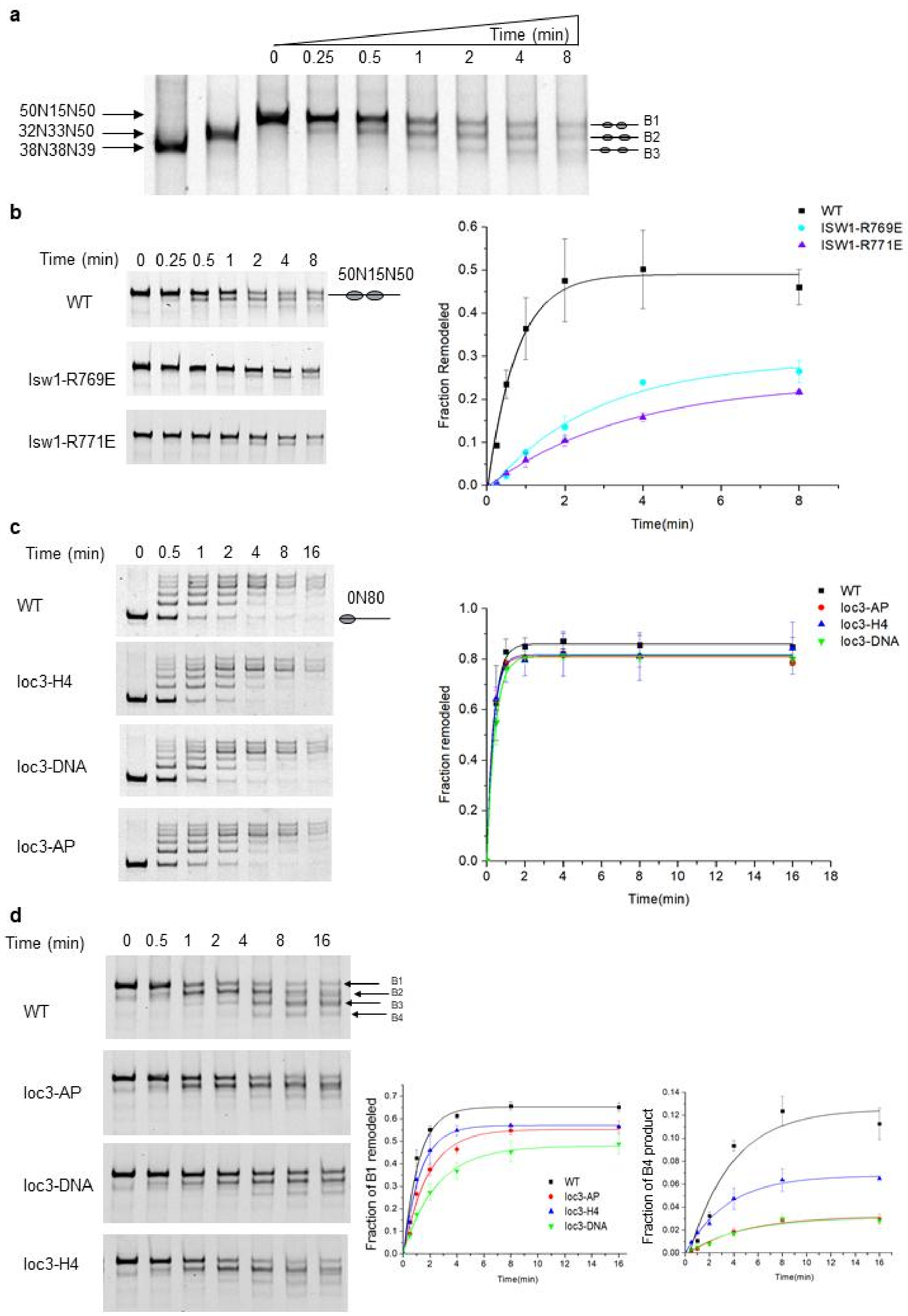


**Supplementary Figure 4 | Additional biochemical analyses of the Isw1a complex.**

(**a**) The ISW1a complex slides the closely packed 50N15N50 dinucleosome to more evenly spaced positions. The dinucleosomes 38N38N39 (evenly spaced nucleosomes, lane 1) and 32N33N50 (one nucleosome moved, lane 2) showed similar migration patterns as those of the ISW1a-remodeling products B2 and B3, respectively. (**b-d**) the chromatin remodeling activities of the WT and indicated mutant ISW1a complex towards the nucleosome substrates 50N15N50 (**b**), 0N80 (**c**), and 10N10N60 (**d**). Representative gels are shown. Quantification of the initial substrates remodeled are shown on the right in (b) and (c); quantification of the initial substrate (B1) and final product (B4) are shown in the middle and right panels of (d), respectively. Error bars indicate SD (n = 3).

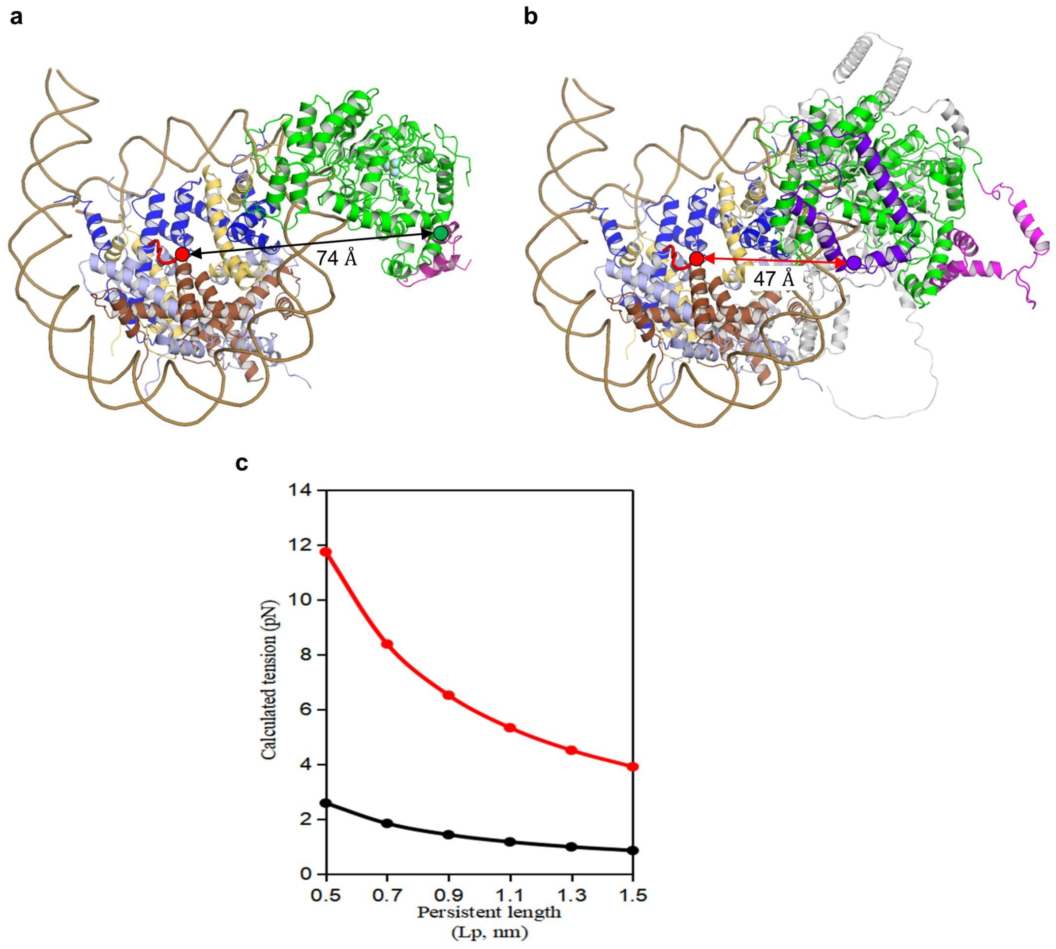


**Supplementary Figure 5 | Tension estimation imposed on NegC of Isw1.** (**a**) The distance between the RA and the motor domain of Isw1 through a disordered NegC domain as reported in current study. The NegC connects the C-terminus of the motor domain with the RA motif, spanning a distance ~74 Å through a disordered sequence of 108 aa (residues 657-764). (**b**) The distance between the RA and the motor domain of Isw1 through the fully folded NegC domain as predicted by AlphaFold (ID: AF-P38144-F1). The distance is ~47 Å through a disordered sequence of 24 aa (residues 741-764). (**c**) Tension estimated by the worm-like-chain model imposed on the disordered sequence of the melted (back line) and folded (red line) NegC. Assuming the persistent length of 1 nm for the polypeptide, the tensions are estimated to be ~1 pN and ~6 pN for the melted and folded NegC, respectively.

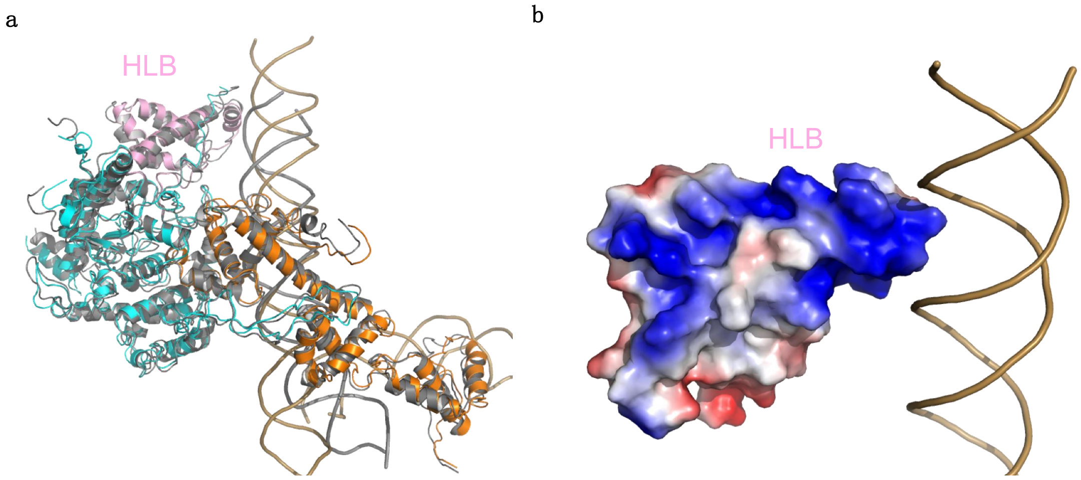


**Supplementary Figure 6 | Interaction between the HLB domain of Ioc3 and the DNA.** (**a**) Structural comparison of the HSS-Ioc3 DNA-binding module bound to the dinucleosome (color coded) and the free DNA (colored grey, PDB code 2Y9Z)[3]. The structures of Ioc3 are aligned. (**b**) Interaction between the HLB domain and the DNA. The electrostatic potential of the HLB domain is calculated by Pymol.

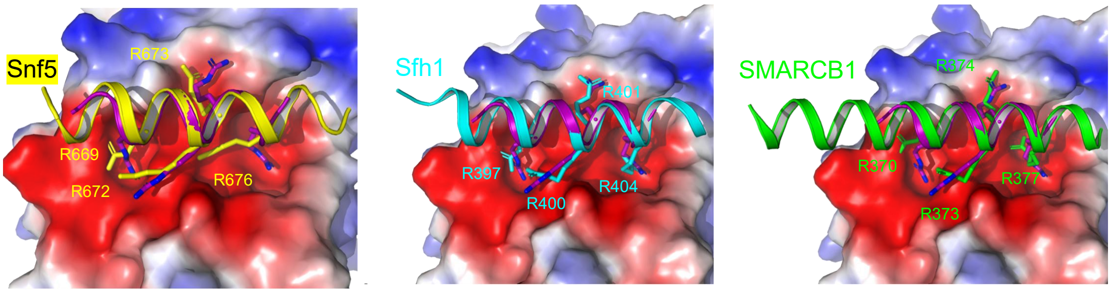


**Supplementary Figure 7 | Comparison of the Finger Helix-nucleosome interaction.** The structures of Finger Helices of Ioc3 (colored violet), Snf5 (PDB code 7egp, colored yellow)[4], Sfh1 (PDB code 5tda, colored cyan)[5], and SMARCB1 (PDB code 6ltj, colored green)[6] are aligned. The electrostatic potentials of the H2A-H2B are calculated by Pymol.

a

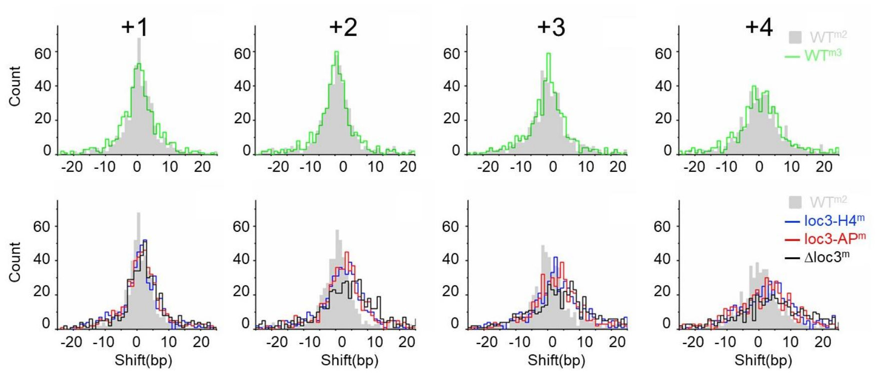


b

| Median Shift(bp) | +1 | +2 | +3 | +4 |
|---|---|---|---|---|
| WT$^{m2}$ | 0 | 0 | -0.5 | 0 |
| WT$^{m3}$ | 0 | 0 | 0 | 0 |
| loc3-H4$^{m}$ | 1 | 2 | 2 | 3 |
| loc3-AP$^{m}$ | 1 | 2 | 2 | 2 |
| Δloc3$^{m}$ | 2 | 4 | 4 | 4 |

c

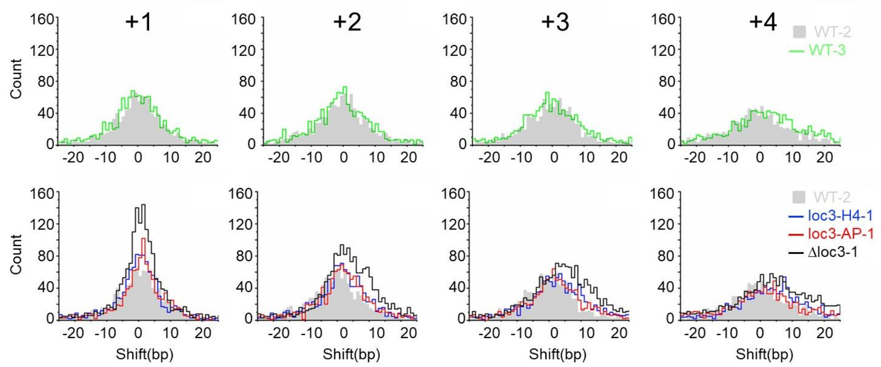


d

| Median Shift(bp) | +1 | +2 | +3 | +4 |
|---|---|---|---|---|
| WT-2 | -1 | -1 | -1 | -1 |
| WT-3 | -1 | -1 | -1 | 1 |
| loc3-H4-1 | 0 | 1 | 3 | 4 |
| loc3-AP-1 | 1 | 1 | 2 | 1 |
| Δloc3-1 | 0 | 3 | 4 | 4 |

e

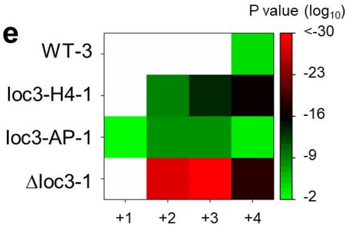


f

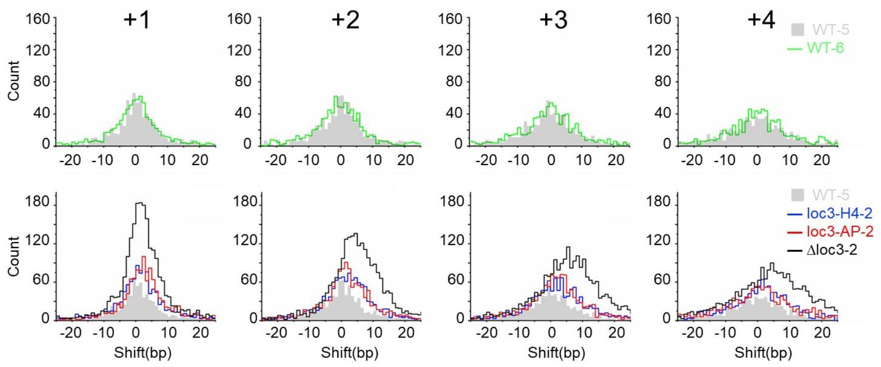


g

| Median Shift(bp) | +1 | +2 | +3 | +4 |
|---|---|---|---|---|
| WT-5 | 0 | 0 | -1 | -1 |
| WT-6 | -1 | -1 | 0 | 0 |
| loc3-H4-2 | 1 | 2 | 1 | 1 |
| loc3-AP-2 | 2 | 1 | 1 | 0 |
| Δloc3-2 | 1 | 4 | 5 | 4 |

h

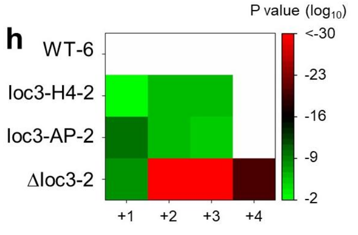


**Supple**

**mentary Figure 8 | Additional analysis of the nucleosome organization of the WT and Ioc3 mutant yeast cells.** (**a-b**) Nucleosome shift analysis of merged data derived from the 542 overlapping genes in the Venn diagram of Figure 5b. To ensure the data quality, each biological replicate dataset includes three WT samples (WT-1, WT-2, and WT-3 in replicate 1; WT-4, WT-5, and WT-6 in replicate 2). WT-1 and WT-3 are merged as the reference; WT-2 and WT-5 are merged as $WT^{m2}$; WT-3 and WT-6 are merged as $WT^{m3}$. (**a**) Histograms of the number of genes having a given nucleosome shift (1-bp bins). (**b**) List of median shifts of the +1 to +4 promoter nucleosomes of the 542 overlapping genes. (**c-e**) Nucleosome shift analysis of biological replicate 1. All genes with at least one significantly shifted promoter nucleosomes are included. (**c**) Histograms of the number of genes having a given nucleosome shift (1-bp bins). (**d**) List of median shifts of the +1 to +4 promoter nucleosomes. The shifts of one of the WT cells are used for Wilcoxon-Mann-Whitney test shown as heat-map in (**e**). (**f-h**) Equivalent plots for biological replicate 2.

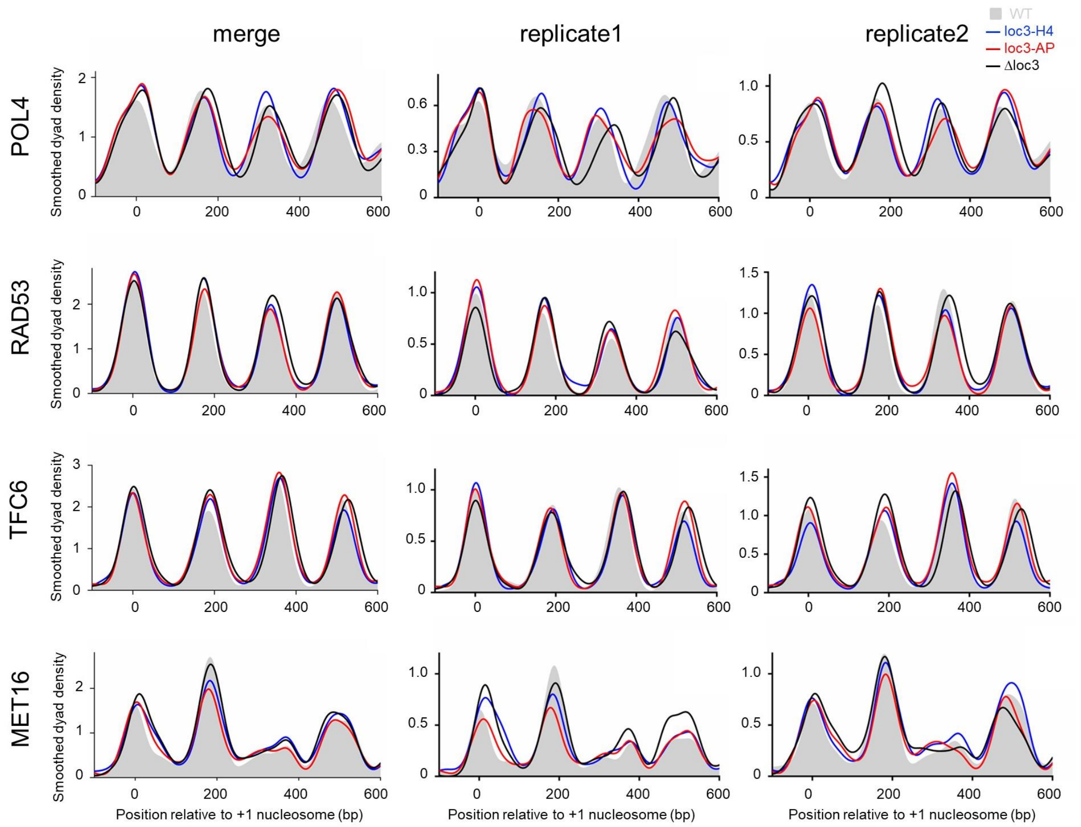


**Supplementary Figure 9 | Four specific loci with shifted nucleosomes.**


1 Schalch, T., Duda, S., Sargent, D. F. & Richmond, T. J. X-ray structure of a tetranucleosome and its implications for the chromatin fibre. *Nature* **436**, 138-141 (2005). https://doi.org:10.1038/nature03686

2 Adhireksan, Z., Sharma, D., Lee, P. L. & Davey, C. A. Near-atomic resolution structures of interdigitated nucleosome fibres. *Nat Commun* **11**, 4747 (2020). https://doi.org:10.1038/s41467-020-18533-2

3 Yamada, K. *et al.* Structure and mechanism of the chromatin remodelling factor ISW1a. *Nature* **472**, 448-453 (2011). https://doi.org:10.1038/nature09947

4 He, Z., Chen, K., Ye, Y. & Chen, Z. Structure of the SWI/SNF complex bound to the nucleosome and insights into the functional modularity. *Cell Discov.* **7**, 28 (2021). https://doi.org:10.1038/s41421-021-00262-5

5 Wagner, F. R. *et al.* Structure of SWI/SNF chromatin remodeller RSC bound to a nucleosome. *Nature* **579**, 448-451 (2020). https://doi.org:10.1038/s41586-020-2088-0

6 He, S. *et al.* Structure of nucleosome-bound human BAF complex. *Science* **367**, 875-881 (2020). https://doi.org:10.1126/science.aaz9761
