## Supplemental Table 1 for "Structure of the ISW1a complex bound to the dinucleosome"

**Supplementary Table 1 Cryo-EM data collection, refinement and validation statistics**

| | #1 ISW1a-diNCP (EMD-32992, PDB 7X3T) | #2 $N_1$-motor (EMD-32995, PDB 7X3W) | #3$N_1$-RA EMD-32996, PDB 7X3X) | #4 $N_2$-Ioc3 (EMD-32994, PDB 7X3V) |
|---|---|---|---|---|
| **Data collection and processing** | | | | |
| Microscope | Krios G3i | Krios G3i | Krios G3i | Krios G3i |
| Camera | K3 | K3 | K3 | K3 |
| Magnification (nominal) | 81000 | 81000 | 81000 | 81000 |
| Electron exposure ($e^{-}$/Å$^2$) | 50 | 50 | 50 | 50 |
| Number of frames collected | 32 | 32 | 32 | 32 |
| Energy filter slit width (eV) | 20 | 20 | 20 | 20 |
| Automation software | AutoEMation2 | AutoEMation2 | AutoEMation2 | AutoEMation2 |
| Voltage (kV) | 300 | 300 | 300 | 300 |
| Micrographs (no.) | 16488 | 16488 | 16488 | 16488 |
| Defocus range (μm) | -1.4— -1.8 | -1.4— -1.8 | -1.4— -1.8 | -1.4— -1.8 |
| Pixel size (Å) | 0.54125 | 0.54125 | 0.54125 | 0.54125 |
| Symmetry imposed | C1 | C1 | C1 | C1 |
| Initial particle images (no.) | 5,769,946 | 5,769,946 | 5,769,946 | 5,769,946 |
| Final particle images (no.) | 66,852 | 69,755 | 170,126 | 212,026 |
| Error of translations | 1.408 | 0.607 | 0.828 | 0.672 |
| Error of rotations | 2.293 | 1.186 | 1.561 | 1.077 |
| Map resolution (Å) (masked) | 5.4 | 3.1 | 3.2 | 3.1 |
| FSC threshold | 0.143 | 0.143 | 0.143 | 0.143 |
| Map sharpening *B* factor (Å$^2$) | -150.9 | -69.4 | -95.2 | -96.2 |
| **Refinement** | | | | |
| Initial model used (PDB code) | | | | |
| Refinement package | | Phenix | Phenix | Phenix |
| Model-map scores | | | | |
| CC(mask) | | 0.87 | 0.87 | 0.82 |
| CC(box) | | 0.86 | 0.89 | 0.84 |
| CC(peaks) | | 0.81 | 0.82 | 0.78 |
| CC(volume) | | 0.87 | 0.86 | 0.84 |
| R.m.s. deviations | | | | |
| Bond lengths (Å) | | 0.011 | 0.010 | 0.011 |
| Bond angles (°) | | 0.774 | 0.814 | 0.761 |
| C-beta deviation | | 0.00 | 0.00 | 0.00 |
| EMRinger score | | 2.93 | 3.57 | 2.97 |
| CaBLAM outliers | | 3.37 | 0.95 | 3.05 |
| **Validation** | | | | |
| MolProbity score | | 1.91 | 1.36 | 1.91 |
| Clashscore | | 10.42 | 6.49 | 9.54 |
| Poor rotamers (%) | | 0.00 | 0.16 | 0.00 |
| Ramachandran plot | | | | |
| Favored (%) | | 94.52 | 98.67 | 93.84 |
| Allowed (%) | | 5.48 | 1.33 | 6.16 |
| Disallowed (%) | | 0.00 | 0.00 | 0.00 |
